# Novel high throughput 3D ECM remodeling assay identifies MEK as key driver of fibrotic fibroblast activity

**DOI:** 10.1101/2024.08.26.609699

**Authors:** Chen-Yi Liao, Jasmijn HM Hundscheid, Justin Crawford, Peter ten Dijke, Beatrice Coorneaert, Erik HJ Danen

**Author notes:** Correspondence to Erik HJ Danen.

## Abstract

In fibrotic tissues, activated fibroblasts remodel the collagen-rich extracellular matrix (ECM). Intervening with this process represents a candidate therapeutic strategy to attenuate disease progression. Models that generate quantitative data on 3D fibroblast-mediated ECM remodeling with the reproducibility and throughput needed for drug testing are lacking. Here, we develop a model that fits this purpose and produces combined quantitative information on drug efficacy and cytotoxicity. We use microinjection robotics to design patterns of fibrillar collagen-embedded fibroblast clusters and apply automated microscopy and image analysis to quantify ECM remodeling between-, and cell viability within clusters of TGFβ-activated primary human skin or lung fibroblasts. We apply this assay to compound screening and reveal actionable targets to suppress fibrotic ECM remodeling. Strikingly, we find that after an initial phase of fibroblast activation by TGFβ, canonical TGFβ signaling is dispensable and, instead, non-canonical activation of MEK-ERK signaling drives ECM remodeling. Moreover, we reveal that higher concentrations of two TGFβ receptor inhibitors while blocking canonical TGFβ signaling, in fact stimulate this MEK-mediated profibrotic ECM remodeling activity.

## INTRODUCTION

Tissue damage, whether caused by trauma, chemical exposure, microbial infection, or genetic defects causing disease, triggers a fibrotic reaction. Fibrosis is essentially a reversible reaction to tissue injury aimed at repair, and its temporal and spatial coordination is crucial for a controlled healing process [1]. Fibrosis is initiated when immune cells that infiltrate the damaged tissue produce cytokines creating an inflammatory environment. This leads to activation of macrophages and the accumulation of myofibroblasts, which originate through activation of resident fibroblasts, though lineage tracing has identified other myofibroblast sources as well [2–4]. One major pathway driving fibrosis involves the production of inflammatory cytokines and transforming growth factor-β (TGFβ) by macrophages, which stimulates the myofibroblasts to produce excessive amounts of collagen-rich extracellular matrix (ECM) [1, 5–7] Changes in proteolytic activity, production of crosslinking enzymes, and high contractile forces that myofibroblasts apply onto the ECM, further lead to extensive remodeling of the healthy tissue matrix into a scar matrix [8, 9]. Fibroblasts typically reside in the stroma as individual cells but have been shown to form networks, communicate through gap junctions, and form clusters in wounded, fibrotic, and cancer tissues [10–14].

As the trigger disappears at the end of the healing process, immune activity is suppressed, fibroblasts return to a quiescent state, and a healthy tissue matrix is restored. However, chronic injury can lead to progressive, irreversible fibrosis and ultimately cause failure of organs such as lung, liver, kidney, heart, skin, and intestinal fibrosis [15–19]. Whereas in each of these cases distinct, organ-specific subsets of immune cells and fibroblast-like cells are involved, the disease in essence follows the same steps in each organ; with sustained immune activity resulting in continuous fibroblast activity and tissue remodeling resulting in loss of proper organ function [20]. Even within a single organ, distinct pathologies with specific clinical features and prognoses can be associated with different underlying triggers causing the fibrotic response [21–23], yet in each case excessive activity of immune and fibroblast cells leading to fibrotic tissue remodeling is involved.

Chronic fibrosis is an important unmet medical challenge that can contribute to organ failure or lack of defense against microbial infections [5]. In many cases the initial cause triggering fibrosis is unknown or might have happened in the past (e.g., cardiac fibrosis due to infarction) making it impossible to target the underlying cause. Rather, drugs that would reduce progressive fibrotic tissue remodeling and tissue stiffening or even cause “tissue normalization” are expected to reduce symptoms and delay or prevent disease progression. Whether advanced fibrosis can be reversed to a normal tissue architecture is controversial but attenuating the ongoing tissue remodeling that causes progressive loss of normal function is expected to benefit many patients.

Current antifibrotic approaches typically target the inflammatory response (the influx of immune cells) but results are not encouraging, and immune cells may also be important for dampening the fibrotic process. Also, clinical translation of candidate drugs targeting the TGFβ signaling axis, are hampered by severe adverse on-target side effects due to the role of TGFβ in maintaining tissue homeostasis [24, 25]. Targeting ⍺vβ6 (an integrin expressed on epithelial cells that supports activation of the ECM-associated latent TGFβ complex such that TGFβ can bind and activate its receptor), is tested in several clinical trials but dosing of antibodies and small molecule drugs requires further optimization to identify a “sweet spot” where clinical benefit outweighs on-target adversity [1, 26, 27]. An attractive alternative would be to target the ECM remodeling capacity of the activated fibroblasts more directly to suppress ongoing tissue remodeling or to even “normalize” the already affected tissue architecture. A recent promising example for this approach is the inhibition of lysyl oxidase-like 2, an ECM crosslinking enzyme. This has been shown to be effective in animal models for melanoma and liver and lung fibrosis [28] and is now in early clinical development for treatment of idiopathic pulmonary fibrosis (http://clinicaltrials.gov/ct2/show/NCT01242189).

Animal models have been used to discover mechanisms underlying for instance lung fibrosis [26]. However, such models lack throughput for larger scale drug testing. Drug screening can be performed on standard 2D “on plastic” cultures using expression of alpha smooth muscle actin (⍺SMA) as a biomarker for myofibroblasts. This approach has also been applied to fibroblasts seeded in 3D collagen gels in multi-well plates and subsequently isolated for measurement of ⍺SMA expression by western blot [29]. However, this does not address actionable mechanisms operating independent of ⍺SMA. To analyze pathological activity of fibroblasts more directly, in vitro models have been developed where fibroblasts are cultured in ECM gels and shrinkage of the collagen gel due to contractive forces generated by the scattered fibroblasts is measured [30]. Yet, this approach is cumbersome, subject to high variation, and difficult to scale up towards a multi-well plate format for drug testing. Lastly, organ-on-chip models have been developed that are highly promising as replacement for animal models to study fibrosis, but throughput of these models is still low, rendering them unfit for larger scale drug testing [31].

Here, we build on our previously established technology [32–34] to develop a method for high-throughput analysis of ECM remodeling between clusters of activated primary human fibroblasts. We apply this assay to compound screening and reveal actionable targets to suppress fibrotic ECM remodeling.

## RESULTS

### Development of a high-throughput model allowing quantitative assessment of fibrotic 3D ECM remodeling

Dermal fibroblasts, isolated from healthy donors, were stimulated with TGFβ, a known profibrotic activator of fibroblast-mediated ECM remodeling [5]. Clusters with a diameter of 200 μm containing ∼500 normal fibroblasts (NF) or TGFβ pre-activated fibroblasts (AF), were printed using automated image guided injection in 96-well plates pre-filled with collagen matrix as we previously described for cancer cells [32]. Subsequently, 3D cultures were exposed to either control medium (CTR), TGFβ, or sphingosine-1-phosphate (S1P) as an alternative profibrotic stimulus [35]. As a first approach, the ability of a single cluster of fibroblasts to align the surrounding collagen fiber network perpendicular to the cluster was analyzed. After confocal reflection microscopy to visualize the 3D ECM fiber network, 4 regions of interest (ROIs) covering 4 quadrants of the image were selected in each Z-section for quantitative image analysis (Fig S1A,B; note the bright spot in each of the 4 stitched images, which is an artefact caused by reflection from optical elements in the microscope [36].

Reflection microscopy images for NF showed minimal changes in response to TGFβ or S1P. For AF, a fiber network was seen in the reflection microscopy images at 24h under CTR and TGFβ conditions (Fig S1C). This did not further increase at later time points for the CTR condition and gradually became somewhat more prominent when exposed to TGFβ. Exposure to S1P appeared to suppress formation of such a fiber network at 24h but subsequently led to the emergence of a very prominent fiber network at 48h and 72h exposure times. This behavior coincided with extensive migration of AF through the collagen network under CTR conditions and slightly less migration in the presence of TGFβ. Exposure to S1P, however, caused an almost complete block of migration (Fig S1C,D). This migration arrest caused by S1P was not observed for NF. Quantitative image analysis using CurveAlign [37] confirmed that AF clusters at 24h were surrounded by a network of aligned fibers in CTR or TGFβ conditions, which was reduced by S1P (Fig S1E). Moreover, the alignment index of the fiber network under S1P conditions strongly increased over time and bypassed that observed under CTR or TGFβ conditions. The same trend was detected when the density of the aligned fiber network was analyzed with CurveAlign (Fig S1F; note the bright spot caused by a reflection artefact is not included in this analysis as CurveAlign only includes aligned fibers).

To simplify the reflection microscopy readout and increase the assay window, we adapted the design by printing multiple fibroblast clusters at 900 μm distance in each well and imaged the ECM network between the clusters (Fig 1A). For NF clusters, refection microscopy showed little change except for a modest increase in signal between clusters exposed to TGFβ for 72 hours (Fig 1B). By contrast, AF clusters exposed to TGFβ or S1P for 72 hours showed a prominent increase in reflection signal between the clusters. We analyzed the total reflective signal intensity applying the same ROI to each Z-section for each condition (Fig 1A,C; note that the bright spot caused by the reflection artefact is excluded from the ROI). This analysis confirmed the modest activation of NF clusters only by TGFβ and detected a strong increase in reflective signal intensity for TGFβ and S1P treated AF clusters, which appeared at an earlier time point for TGFβ as compared to S1P (Fig 1C,D).

**Figure 1.**
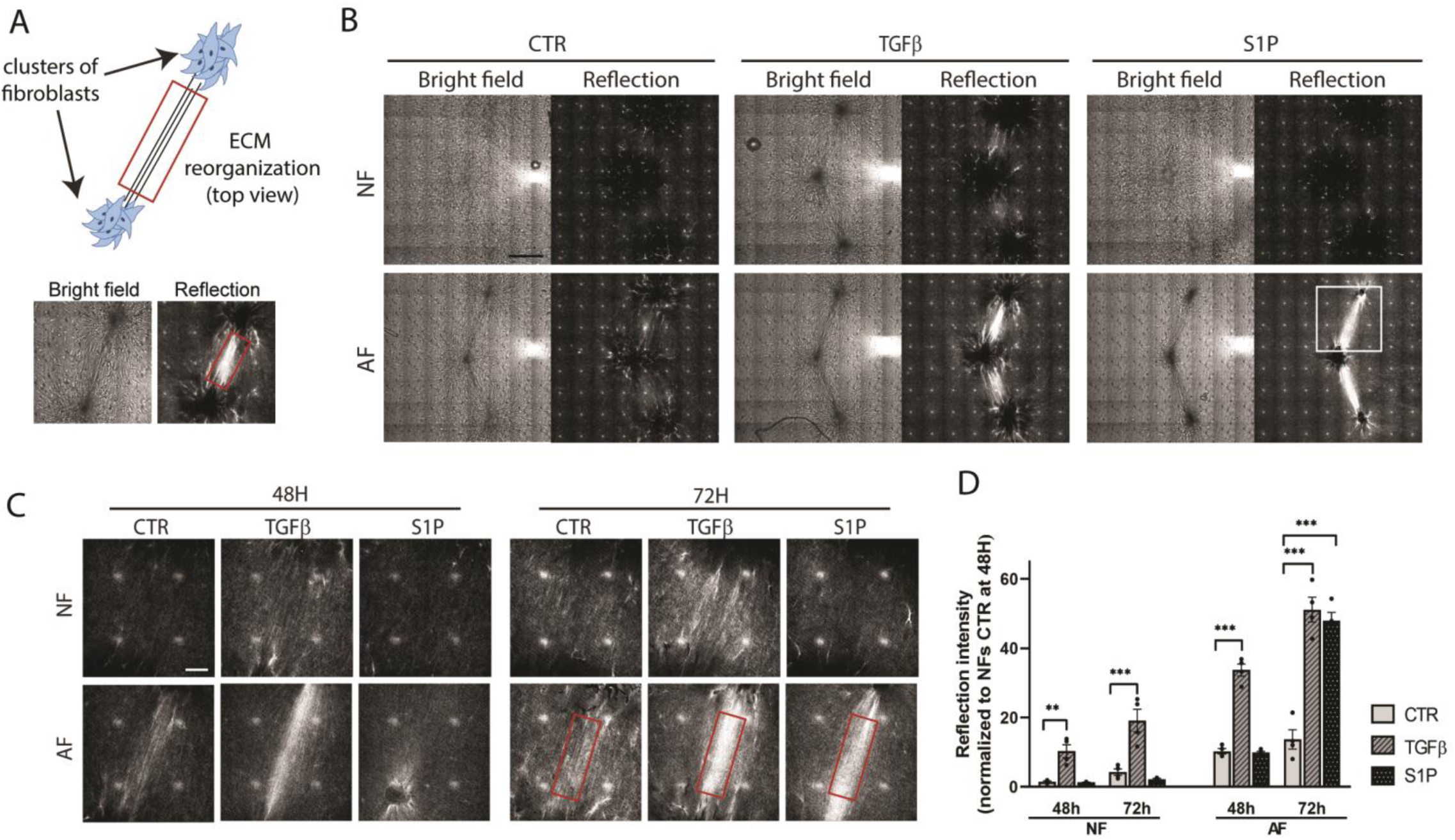
Multi-cluster model for quantitative assessment of TGFβ- and S1P-induced ECM remodeling. **(A)** Cartoon showing model design and brightfield and reflection images of the ROI after 72 hours TGFβ treatment. **(B)** Multiple clusters of NF or AF printed in collagen at 900 μm distance and maintained in CTR medium or stimulated with TGFβ or S1P for 72 hours. Brightfield and reflection images are shown of a single Z-section through the clusters. Note that the pattern of white spots in the reflection images is an artefact caused by reflection from optical elements in the microscope [36]. Bar = 500 μm. The white square indicates the area depicted in (C). **(C)** Single Z-section reflection images for NF and AF collagen-embedded fibroblast clusters treated with or without TGFβ or S1P for 48 and 72 hours. Note that the ROI (red rectangle) does not include the white dots caused by the reflection artefact. Bar = 100 μm. **(D)** Quantification of reflection signal from (C) taking the average reflection signal density in the ROI across the Z-stack. Graphs show the mean and SEM of two independent experiments, each performed in duplicate. **, p < 0.01; ***, p < 0.001. NF; normal fibroblasts; AF, TGFβ-activated fibroblasts.

Taken together, this data indicates that both pre-treatment and prolonged exposure to profibrotic triggers in the 3D culture affect migration and ECM remodeling by ECM embedded clusters of fibroblasts. Exposure to S1P blocks fibroblast migration and strongly stimulates ECM remodeling but only when fibroblasts are pre-activated with TGFβ. Exposure to TGFβ leads to a more gradual increase in ECM remodeling that is mild in NF and more robust in AF and most prominent in the multi-cluster model.

### Mechanistic underpinning of the confocal reflection microscopy density readout

For further experiments we selected the multi-cluster model that provided a satisfactory assay window, reproducibility, and throughput for testing of candidate anti-fibrotic drugs. We first analyzed the changes induced by TGFβ that could contribute to the confocal reflection intensity readout parameter. Antibody-mediated collagen staining confirmed that regions with increased intensity in the reflection microscopy were enriched in collagen (Fig 2A). The conversion of normal fibroblasts to myofibroblasts is associated with an upregulation of the expression of ⍺SMA and fibronectin (FN) [38–40]. An increase in FN staining was observed between the clusters of AF as compared to NF (Fig 2B). No obvious increase was observed in FN expression within the fibroblast clusters, indicating that FN was rapidly secreted into the ECM network (Fig 2C). Under TGFβ stimulation, at the interface between the cells and the ECM, FN formed elongated fibrils that connected to F-actin filaments within the cells, something that was not observed under control conditions (Fig 2D). Furthermore, under TGFβ stimulation, F-actin stress fibers in fibroblasts in contact with the ECM bridging the clusters, showed parallel alignment while F-actin in unstimulated fibroblast clusters appeared less organized (Fig 2E,F). Lastly, TGFβ treatment of AF did not increase ⍺SMA expression but reduced fibroblast migration in the presence of TGFβ caused ⍺SMA staining to be more concentrated in the core of the fibroblast clusters as compared to CTR conditions (Fig 2G).

**Figure 2.**
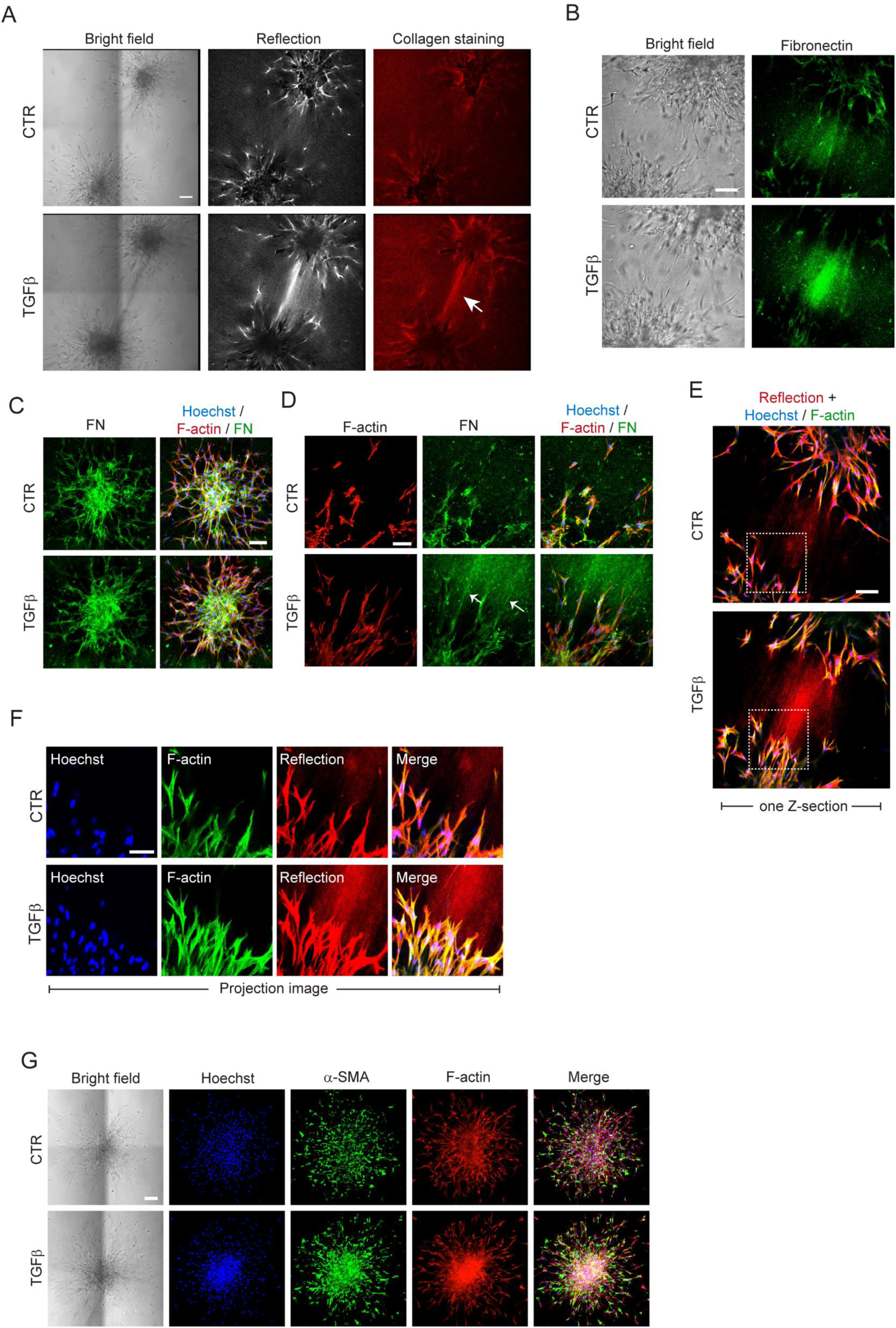
TGFβ stimulated fibroblast clusters show cells with prominent F-actin fibers interacting with a bundle of remodeled ECM containing collagen and FN. **(A)** Brightfield, reflection, and collagen immunostaining of collagen-embedded AF clusters exposed to vehicle control (CTR) or TGFβ containing media. Arrow indicates collagen bundle formed between two fibroblast clusters in presence of TGFβ. The image shows a single Z-section through the clusters. Bar = 100 μm. **(B)** Brightfield and FN immunostaining of collagen-embedded AF clusters exposed to CTR or TGFβ containing media. The image shows a single Z-section through the clusters. Bar = 100 μm. **(C)** Maximum projection images showing FN expression within fibroblast clusters in CTR or TGFβ containing media. Bar = 100 μm. Blue, Hoechst; Red, F-actin; Green, FN. **(D)** F-actin and FN staining shows F-actin fibers in protruding fibroblasts connecting to FN bundles (white arrows) under TGFβ condition. The image shows a single Z-section through the clusters. Blue, Hoechst; Red, F-actin; Green, FN. Bar = 100 μm. **(E)** F-actin staining and reflection microscopy shows protruding fibroblasts connecting to the bridge of remodeled ECM between the clusters under TGFβ condition. The image shows a single Z-section through the clusters. White box indicates the area shown in (F). Bar = 100 μm. **(F)** Maximum projection images showing F-actin fibers in protruding fibroblasts connecting to the bridge of remodeled ECM between the clusters in TGFβ exposed conditions. **(G)** Brightfield and maximum projection images showing ⍺SMA and F-actin expression in AF cluster exposed to CTR or TGFβ containing media. Blue, Hoechst; Green, ⍺SMA; red, F-actin. Bar = 100 μm.

### Application of the model to drug testing in primary dermal and lung fibroblasts and demonstration that drugs inhibiting cytoskeletal contractility or collagen cross-linking attenuate ECM remodeling

We explored application of the developed model in testing the efficacy of candidate anti-fibrotic compounds. In addition to excessive production of ECM, fibrotic ECM remodeling involves increased application of contractile forces onto the ECM fiber network by fibroblasts and enhanced ECM crosslinking [41, 42]. We first focused on the pathway that supports cell contractility. Rho-associated protein kinase (ROCK) stimulates myosin light chain activity to promote actomyosin contractility. It also activates LIN-11, Isl-1, MEC-3 kinase (LIMK), which, in turn, phosphorylates cofilin to increase F-actin stabilization [41]. Two ROCK inhibitors (ROCKi) and two LIMKi were selected and these suppressed TGFβ-induced ECM remodeling in a dose-dependent manner (Fig 3A,B). Analyzing a complete Z-stack showed that ROCK inhibition led to a full 3D loss of the entire ECM bridge between the two fibroblast clusters (Fig 3C). In parallel to the analysis of ECM remodeling between fibroblast clusters, we used propidium iodide (PI) staining and fluorescence confocal microscopy to monitor loss of viability within the clusters. Indeed, at higher concentrations of the ROCKi, signs of cytotoxicity could be observed (Fig 3D). Quantitative image analysis identified concentrations where ROCKi were effective but not toxic and higher concentrations where toxicity became obvious (Fig 3E).

**Figure 3.**
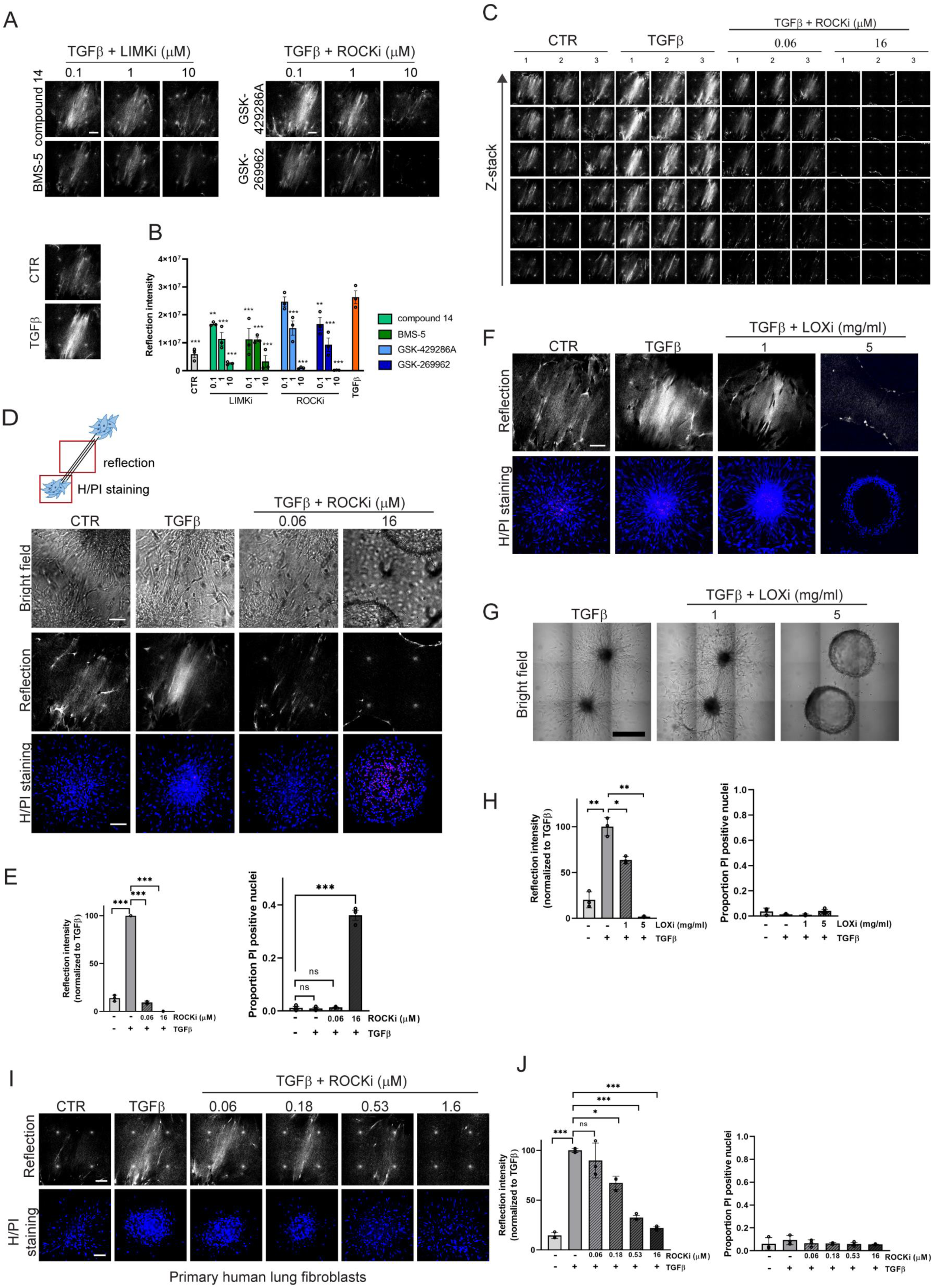
Cytoskeletal contractility and collagen fiber cross-linking are required for TGFβ mediated ECM remodeling by fibroblast clusters and application of the model to lung fibroblasts. **(A)** ECM remodeling visualized by reflection microscopy between two AF clusters stimulated for 72 hours with TGFβ in presence of the indicated concentrations of two LIMKi or ROCKi. CTR and TGFβ conditions without inhibitors are shown below. Bar = 100 μm. **(B)** Quantification of the reflection signals as shown in (A). Graphs show the mean and SEM of two independent experiments, each performed in triplicate. **, p < 0.01; ***, p < 0.001 compared to the TGFβ alone condition. **(C)** Z-stack of reflection images showing ECM remodeling between two AF clusters under CTR or TGFβ stimulation in absence or presence of the indicated concentrations of the GSK-429286A ROCKi. **(D)** Cartoon illustrating imaging areas for assessing ECM remodeling and cell viability (top) and brightfield and reflection (single Z-section) images of the area between AF clusters and Hoechst (blue)/PI (red) staining (maximum projection) inside AF clusters exposed to CTR or TGFβ containing media in absence or presence of the indicated concentrations of the GSK-429286A ROCKi (bottom). Bar = 100 μm. **(E)** Quantification of reflection signals and Hoechst/PI staining as shown in (D). Graphs show the mean and SEM of two independent experiments, each performed in triplicate. ns, non-significant; ***, p < 0.001. **(F)** Reflection (single Z-section) images of the area between AF clusters and Hoechst (blue)/PI (red) staining (maximum projection) inside AF clusters exposed to CTR or TGFβ containing media in absence or presence of the indicated concentrations of the βAPN LOXi. Bar = 100 μm. **(G)** Brightfield images showing collagen-embedded AF clusters exposed for 72 hours to TGFβ in absence or presence of the indicated concentrations of the βAPN LOXi. Bar = 100 μm. **(H)** Quantification of reflection signals and Hoechst/PI staining as shown in (F). Graphs show the mean and SEM of two independent experiments, each performed in triplicate. *, p < 0.05; **, p < 0.01. **(I)** Reflection (single Z-section) images of the area between clusters of activated primary human lung fibroblasts and Hoechst (blue)/PI (red) staining (maximum projection) inside these clusters exposed to CTR or TGFβ containing media in absence or presence of the indicated concentrations of the GSK-429286A ROCKi (bottom). Bar = 100 μm. **(J)** Quantification of reflection signals and Hoechst/PI staining as shown in (I). Graphs show the mean and SEM of two independent experiments, each performed in triplicate. ns, non-significant; *, p < 0.05; ***, p < 0.001.

We next focused on the lysyl oxidase (LOX) enzymes that mediate the formation of intra- and inter-molecular cross-links in collagen molecules within the ECM and have been implicated in the development of fibrosis in various organs [42]. We tested whether an irreversible broad LOX inhibitor (LOXi) 3-Aminopropionitrile fumarate salt (βAPN) [43] attenuated ECM remodeling in our model. This compound led to a reduction in reflection signal density at 1 mg/ml and a virtually complete loss at 5 mg/ml (Fig 3F). Notably, when cultures were treated with 5 mg/ml βAPN, fibroblast migration was suppressed and the shape of the cell cluster transitioned to a hollow sphere even though βAPN did not cause any loss of viability at the concentrations tested (Fig 3G,H).

To demonstrate broader applicability of the assay, we used the same setup as applied to human dermal fibroblasts for the analysis of ECM remodeling by activated primary normal lung fibroblasts. A similar behavior as observed with dermal fibroblasts was seen in this model, with extensive ECM remodeling in response to TGFβ. Moreover, this model was similarly applicable to drug testing and showed that ECM remodeling was dependent on ROCK activity although these cells were less sensitive to cytotoxicity in the presence of increased concentrations of the ROCKi (Fig 3I,J).

Together, this data demonstrates that the developed assay is applicable to primary dermal and lung fibroblasts. The assay allows testing of the effect of candidate drugs on fibrotic ECM remodeling versus their cytotoxicity. It identifies contractility and ECM crosslinking as actionable targets to prevent ECM remodeling.

### Drugs inhibiting cytoskeletal contractility or collagen cross-linking enzymes can attenuate but cannot reverse ECM remodeling by dermal fibroblasts

Having established that inhibition of the enzymes mediating fibroblast contractility or collagen crosslinking could attenuate TGFβ-induced ECM remodeling, we next asked if such treatment could reverse an already established remodeled ECM. AF were printed in a collagen matrix and stimulated by TGFβ for 72 hours to form an ECM bridge between two fibroblast clusters. Subsequently, TGFβ containing medium was replaced with CTR medium with or without a LOXi or a ROCKi and cultures were monitored for an additional 72 hours. Such prolonged treatment with a high concentration of LOXi led to a significant increase in cell death while this was not observed for the ROCKi (Fig 4A,B). Importantly, treatment of fibroblast clusters with concentrations of the LOXi or ROCKi that could prevent ECM remodeling (Fig 3) failed to affect the ECM bridge that was already generated between the clusters (Fig 4A,B). This indicates that a remodeled ECM, once established is largely irreversible.

**Figure 4.**
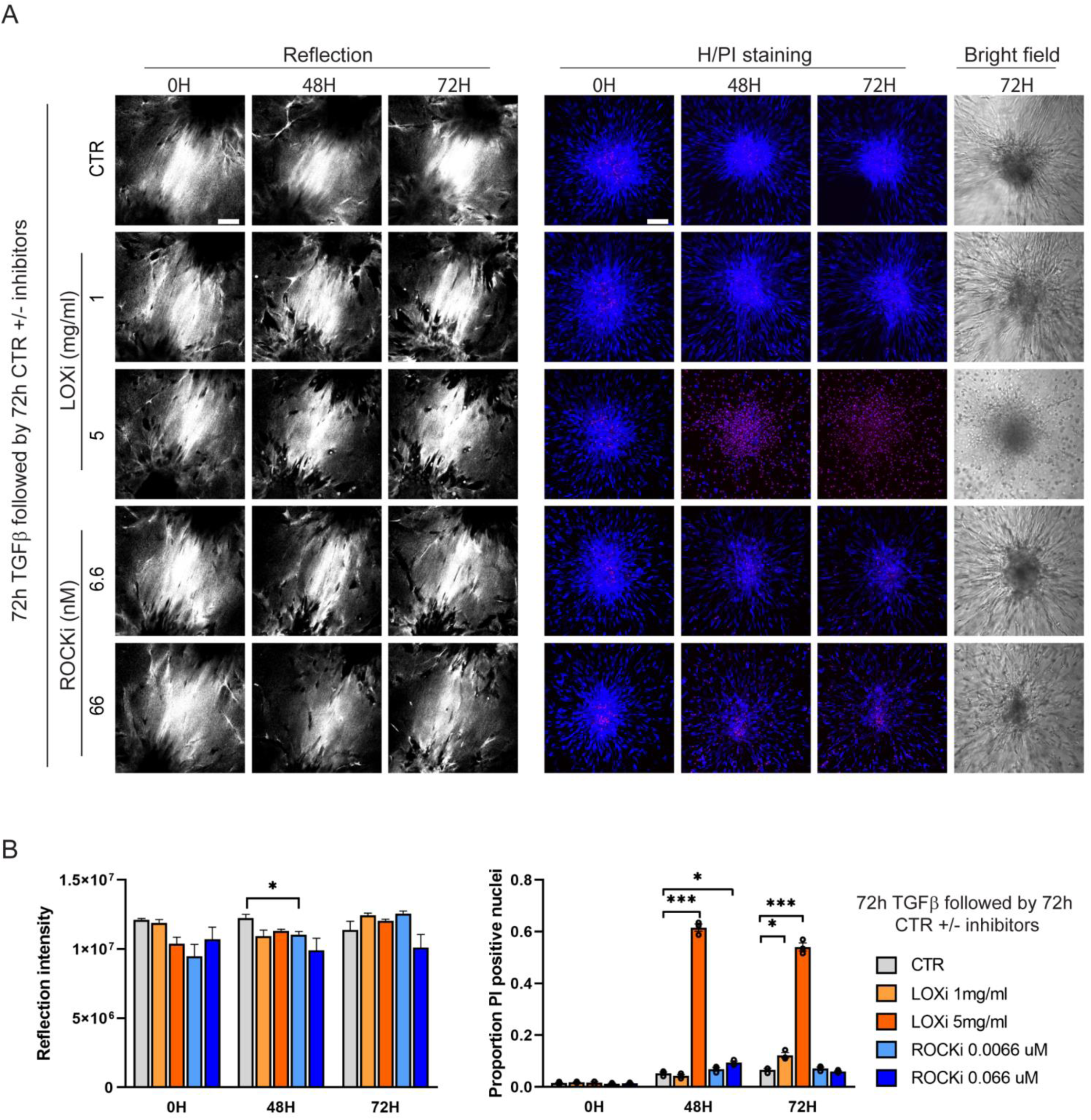
TGFβ mediated ECM remodeling by AF clusters is irreversible. **(A)** Reflection (single Z-section) images of the area between AF clusters and Hoechst (blue)/PI (red) staining (maximum projection) and brightfield images inside AF clusters exposed to TGFβ for 72 hours followed by exposure to CTR media in absence or presence of the indicated concentrations of the GSK-429286A ROCKi or the βAPN LOXi. Bar = 100 μm. **(B)** Quantification of reflection signals and Hoechst/PI staining as shown in (A). Graphs show the mean and SEM of two independent experiments, each performed in triplicate. *, p < 0.05; ***, p < 0.001.

### High-throughput screening for efficacy versus cytotoxicity of candidate anti-fibrotic compounds

Based on the results obtained so far, we designed a screening approach where concentration ranges of candidate anti-fibrotic compounds could be tested to identify a potential window of efficacy without cytotoxicity (Fig 5A). IC50 values in cell-based assays are typically considerably higher than those in cell free, biochemical assays. Concentration ranges we chose started from ∼1000 x IC50_(cell free assay)_ and reached ∼4 x IC50_(cell free assay)_ in 5 steps of 1:3 dilution. For some compounds the maximum concentration was <1000 x IC50_(cell free assay)_ as 20 µM was the highest dose tested to keep the DMSO solvent concentration below 0.2% to prevent DMSO-induced cytotoxicity (Fig 5B).

**Figure 5.**
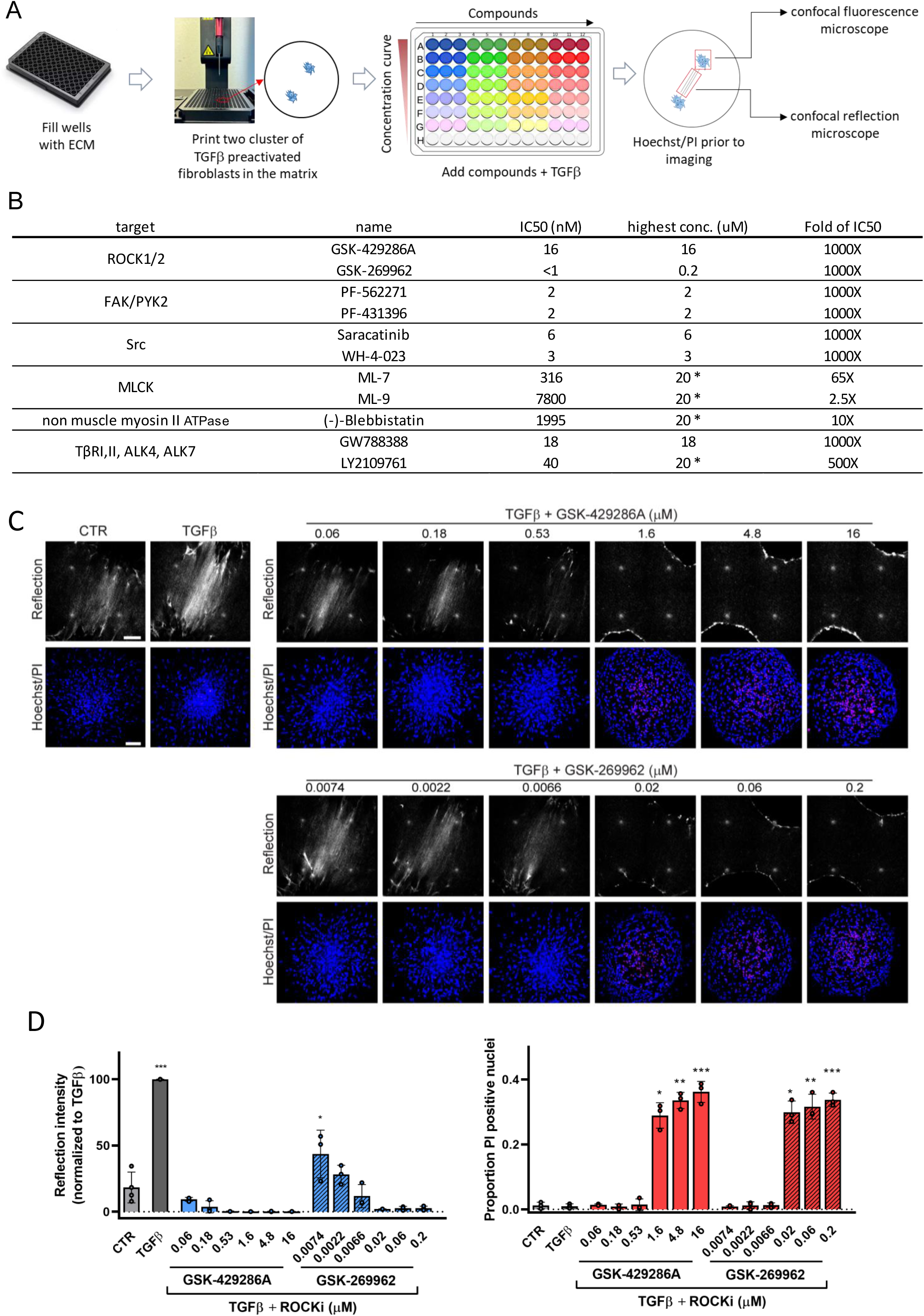
Application of the developed model to screening of candidate anti-fibrotic compounds – ROCKi as example. **(A)** Cartoon showing setup for screening of candidate anti-fibrotic drugs. **(B)** Compounds used in this study with their highest concentration tested based on 1000 x IC50_(cell free assay)_ where possible. *Indicates compounds where 1000 x IC50_(cell free assay)_ could not be reached and a maximum concentration of 20 μM was used to avoid final DMSO solvent concentrations >0.02%. **(C)** Reflection (single Z-section) images of the area between AF clusters and Hoechst (blue)/PI (red) staining (maximum projection) inside AF clusters exposed for 72 hours to CTR or TGFβ containing media in absence or presence of the indicated concentrations of the indicated ROCKi. Bar = 100 μm. **(D)** Quantification of reflection signals and Hoechst/PI staining as shown in (C). Graphs show the mean and SEM of two independent experiments, each performed in triplicate. *, p < 0.05; **, p < 0.01; ***, p < 0.001, compared to CTR.

The screen included two ROCKi, GSK-429266A and GSK-269962. Both inhibitors showed cytotoxicity at the highest three concentrations but appeared effective at lower, non-toxic concentrations (Fig 5C). Quantitative image analysis of data obtained from two independent experiments, each performed in triplicate confirmed that both compounds showed a window where ECM remodeling was suppressed without signs of cytotoxicity (Fig 5D). The series of compounds included in the screen inhibited key regulators of cell contractility (ROCK1/2, non-muscle myosin II ATPase, MLCK), cell-matrix adhesion signaling (FAK, Src), and TGFβ signaling (TGFβ type I receptor kinase (TβRI)/ activin receptor-like kinase 5 (ALK5)) (Fig 5B; Table 1) [41, 44–47]. Most of the compounds showed a dose response with increasing concentrations suppressing ECM remodeling but several of these were only effective in a range where cytotoxicity was prominent, including WH-4-023, ML-7, and ML-9 (Fig 6A; Fig S2).

**Figure 6.**
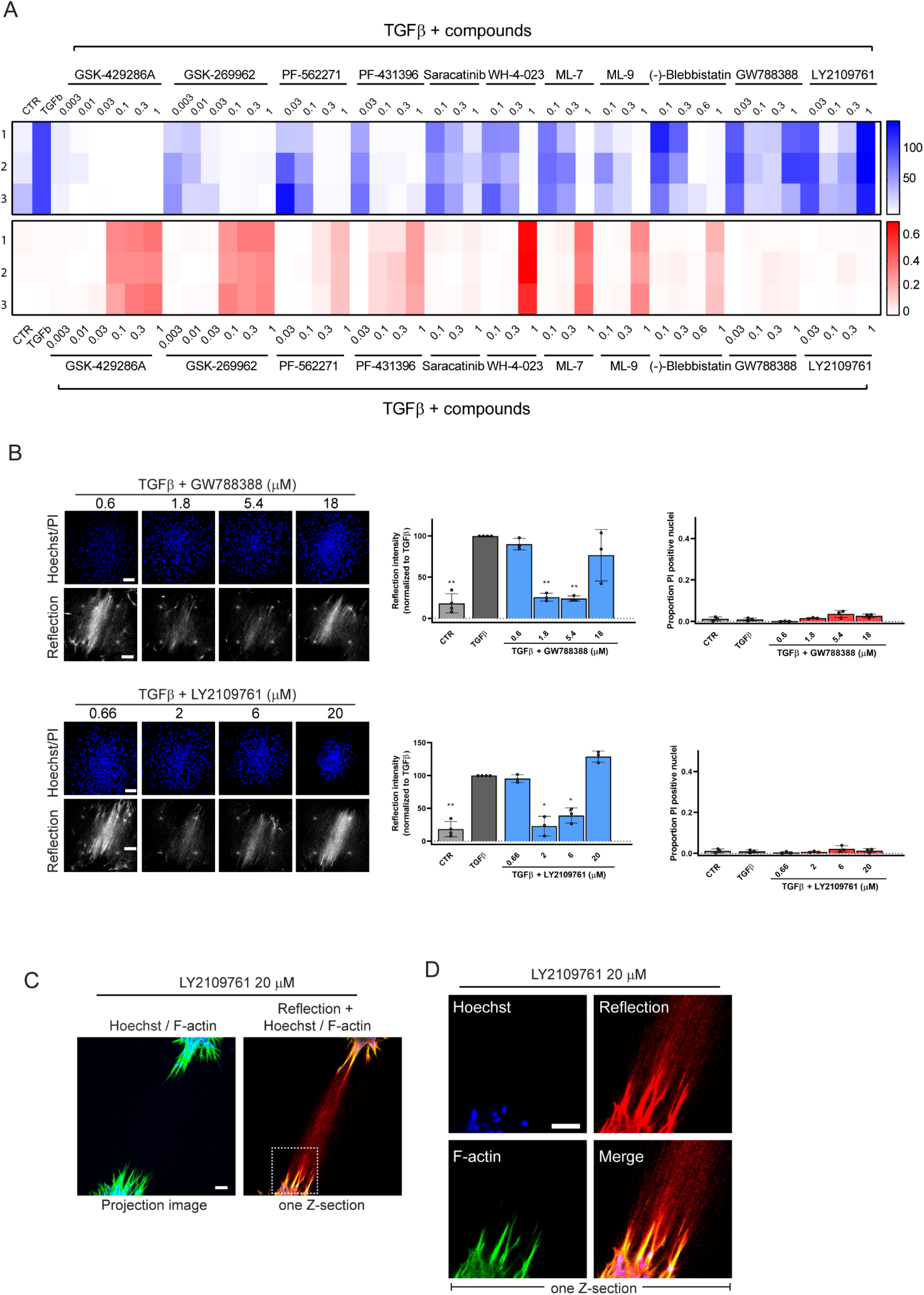
Application of the developed model to screening of candidate anti-fibrotic compounds – unanticipated effect of TβRi. **(A)** Heatmap showing the effect of the indicated concentration ranges of compounds on the reflection signal between TGFβ-stimulated AF cluster (blue; scale ranging from 0 to 150; higher values indicate higher ECM remodeling) and on the proportion of PI-positive cells within the clusters (red; scale ranging from 0 to 1; higher values correspond to higher cytotoxicity). Highest concentration for each compound indicated as ’1’, followed by subsequent 1:3 dilutions. Three replicates are shown vertically. **(B)** Reflection (single Z-section) images of the area between AF clusters and Hoechst (blue)/PI (red) staining (maximum projection) inside AF clusters exposed for 72 hours to TGFβ containing media in absence or presence of the indicated concentrations of the two indicated TβRi. Bar = 100 μm. Graphs show quantification of reflection signals and Hoechst/PI staining. Mean and SEM of two independent experiments, each performed in triplicate is shown. *, p < 0.05; **, p < 0.01 compared to TGFβ alone condition. **(C,D)** Combined Hoechst (blue), F-actin (green), and reflection (red) images of the area between AF clusters treated for 72 hours with CTR medium containing 20 μM of the LY2109761 TβRi. White box in (C) is enlarged in (D). Images are a single Z-section through the clusters. Bar = 100 μm.

**Table 1.** The list of pharmacological inhibitors used in this study.

| Name | Target | Catalog No. | Company |
| --- | --- | --- | --- |
| PF-562271 | FAK/PYK2 | S2890 | SelleckChem |
| PF-431396 | FAK/PYK2 | PF431396 | Charnwood Molecular Ltd |
| GSK-269962 | ROCK1/2 | 1167 | Axon MedChem |
| GSK-429286A | ROCK1/2 | HY-11000 | MedChem Express |
| Saracatinib | Src | S1006 | SelleckChem |
| WH-4-023 | Src | HY-12299 | MedChem Express |
| ML-7 | MLCK | 475880 | Calbiochem |
| ML-9 | MLCK | 475882 | Calbiochem |
| (-)-Blebbistatin | non muscle myosin II ATPase | HY-13441 | MedChem Express |
| GW788388 | TβRI/ALK5, TβRII, ALK4, ALK7 | S2750 | SelleckChem |
| LY2109761 | TβRI/ALK5, ALK4, ALK7, and TβRII | HY-12075 | Haoyuan ChemExpress |
| BMS-5 (LIMKi 3) | LIMK | DSCR001763 | DSK |
| LIMK-IN-1 (Compound 14) | LIMK | EN300-115361 | Enamine |
| 3-Aminopropionitrile fumarate salt | Lysyl oxidase (LOX) | A3134 | Sigma-Aldrich |

The two compounds GW788388 and LY2109761, targeting TβRI (as well as TβRII, ALK4, and ALK7 [48, 49]) showed a very different pattern. Below 1 µM they were not effective, in the low µM range they suppressed ECM remodeling, but at >10 µM they did not appear to affect TGFβ-induced ECM remodeling (Fig 6A,B). Neither of these inhibitors gave rise to cytotoxicity at the concentrations tested. Strikingly, 3D cultures where AF were exposed to such high concentration of LY2109761 in absence of TGFβ, showed fibroblasts protruding from the clusters interacting with an ECM bridge in a manner identical to AF exposed to TGFβ (Fig 6C,D).

This data demonstrates that the developed platform is suitable to high-throughput, high-content screening to assess the efficacy versus cytotoxicity of candidate anti-fibrotic compounds. It identifies candidate actionable targets to inhibit fibrotic ECM remodeling. It shows that for some compounds apparent efficacy in fact overlaps with cytotoxicity. And it indicates that the TβRi, GW788388 and LY2109761, at high concentrations stimulate rather than inhibit ECM remodeling.

### MEK activation is critical for ECM remodeling induced by TGFβ or high concentration TβRi

In the canonical TGFβ pathway, TGFβ binding leads to the formation of a tetrameric complex of TβRI and TβRII and activation of TβRI kinase activity. TβRI subsequently activates SMAD complexes that translocate to the nucleus and activate genes containing a 5’-CAGA-3’ DNA box [50, 51]. To investigate whether high concentrations of a TβRi might unintentionally “turn on” canonical TGFβ signaling, a CAGA_12_-dynGFP reporter was expressed in the fibroblasts. In 2D culture, CAGA_12_-dynGFP expression was strongly upregulated in the presence of TGFβ and 2 or 20 µM LY2109761 blocked this response as expected (Fig 7A). Similarly, in 3D cultures CAGA_12_-dynGFP expression induced by exposure to TGFβ was blocked by both LY2109761 concentrations (Fig 7B,C). However, while 2 µM LY2109761 also suppressed ECM remodeling, 20 µM LY2109761 did not, and this high concentration was able to induce ECM remodeling in absence of TGFβ without inducing canonical TGFβ signaling. ECM remodeling in the presence of 20 µM LY2109761 was blocked by co-treatment with a ROCKi, indicating that a similar, contractility-dependent mechanism as triggered by TGFβ was at play (Fig 7B,C).

**Figure 7.**
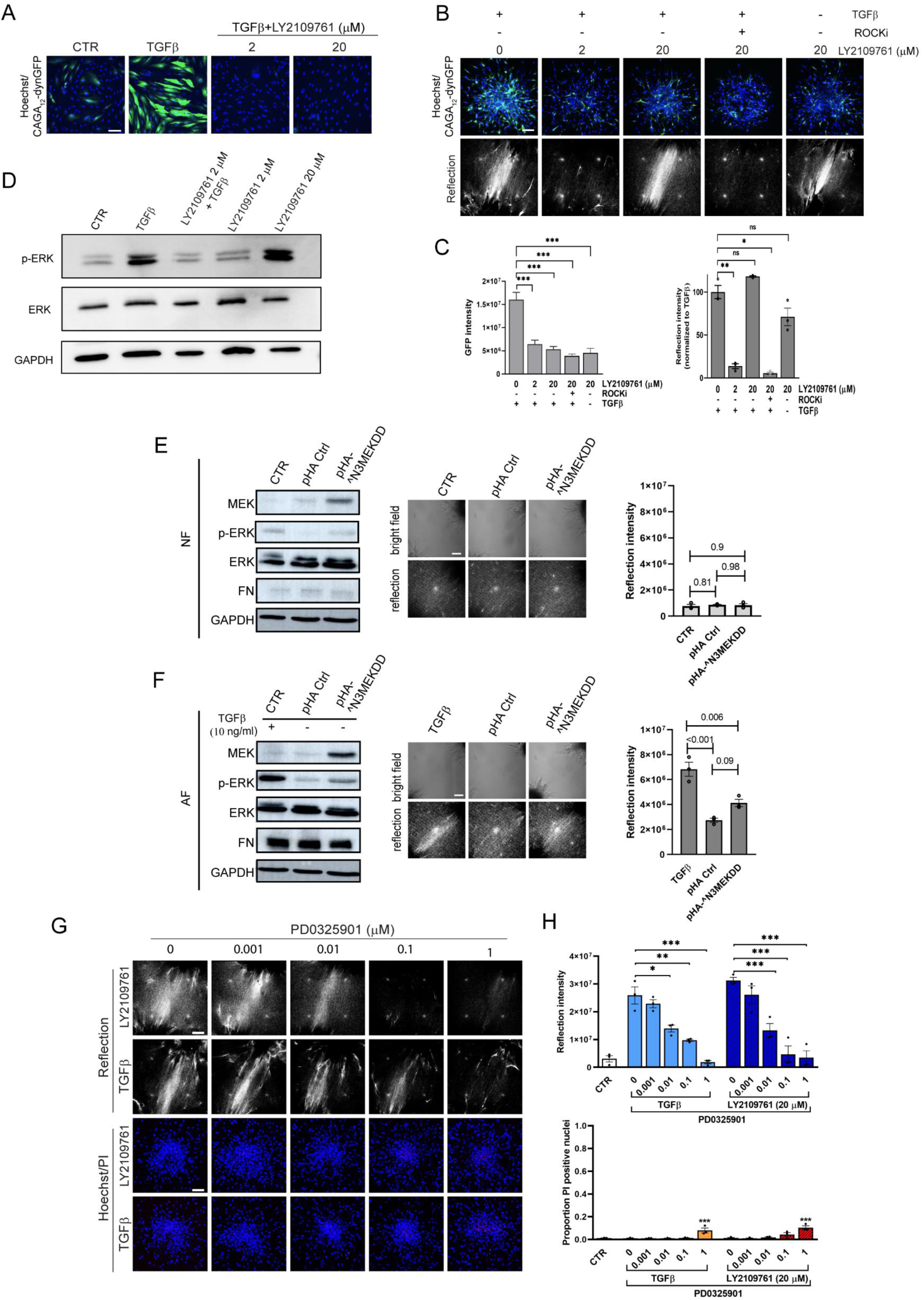
MEK activation is essential for ECM remodeling induced by TGFβ or by high concentration TβRi. **(A)** CAGA_12_-dynGFP signal in 2D cultures of NF treated with CTR medium or TGFβ for 72 hours in absence or presence of the indicated concentrations of the LY2109761 TβRi. Blue, nucleus; Green, CAGA_12_-dynGFP. Bar = 100 μm. **(B)** Reflection (single Z-section) images of the area between AF clusters (bottom) and Hoechst (blue)/ CAGA_12_-dynGFP (green) staining (maximum projection) inside AF clusters exposed for 72 hours to the indicated combinations of TGFβ and compounds. Bar = 100 μm. **(C)** Quantification of reflection signals and GFP intensity from (B). Graphs show the mean and SEM of two independent experiments, each performed in triplicate. ns, not significant; *, p < 0.05; **, p < 0.01; ***, p < 0.001. **(D)** Western blot analysis of total- and phospho-ERK in 2D cultures of AF treated with CTR or TGFβ-containing medium in presence or absence of the indicated concentrations of the LY2109761 TβRi. GAPDH serves as loading control. **(E,F)** NF (E) and AF (F) were transiently transfected with pHA-MEK^N3;S218E;S222D, empty pHA vector, or untransfected, followed by 2 days recovery in control medium. For WB analysis (left panel) of the indicated (phospho)proteins cells were cultured in 2D for 3 days in control medium (E) or medium with or without TGFβ as indicated (F). For brightfield and reflection microscopy for ECM remodeling capacity (right panels), cells were printed to form 3D collagen embedded clusters and exposed for 3 days to the same media. Graphs show mean and SEM of two independent experiments each performed in triplicate. ns, not significant; **, p < 0.01. **(G)** Reflection (single Z-section) images of the area between AF clusters and Hoechst (blue)/PI (red) staining (maximum projection) inside AF clusters exposed for 72 hours to TGFβ or 20 μM LY2109761 in absence or presence of the indicated concentrations of the PD0325901 MEKi. Bar = 100 μm. **(H)** Quantification of reflection signals and Hoechst/PI staining as shown in (G). Graphs show the mean and SEM of two independent experiments, each performed in triplicate. *, p < 0.05; **, p < 0.01; ***, p < 0.001.

Besides canonical SMAD-mediated TGFβ signaling, TGFβ can modulate other signaling pathways, including the activation of Ras-mitogen-activated protein kinase (MAPK) signaling [24, 52]. Indeed, phosphorylation of ERK MAPK in AF was stimulated by TGFβ and this was prevented by exposure to 2 µM LY2109761, whereas exposure to 20 µM LY2109761 mimicked the effect of TGFβ, causing a similar level of ERK phosphorylation (Fig 7D).

To test whether enhanced ERK MAPK activity was sufficient to trigger ECM remodeling, we attempted to express an activated MEK mutant. However, in line with earlier reports showing a senescence response [53], constitutive activation of ERK MAPK signaling in primary fibroblasts was poorly tolerated. Lentiviral expression of a constitutive active MEK mutant led to a loss of almost all cells (while a control GFP-actin virus was well-expressed) (Fig S3). Switching to transient transfection of a CMV-driven constitutive active MEK^N3;S218E;S222D construct did lead to a slight increase in ECM remodeling in AF (but not in NF) as compared to cells expressing a control plasmid, but this did not reach significance (Fig 7E,F). Such inefficiency could be explained by insufficient expression of activated MEK, since the level of p-ERK in the transfected cell pool was low compared to that caused by exposure to TGFβ (Fig 7E,F). Expression of FN, a target of canonical TGFβ signaling, which was low in NF and high in AF, was not influenced by the weak expression of activated MEK (Fig 7E,F). Notably, as TGFβ was absent during the transfection procedure and subsequent culture of the transfected cells, this indicated that TGFβ-mediated conversion to AF is maintained at least 5 days.

To assess whether enhanced ERK MAPK activity was required for ECM remodeling triggered by TGFβ or high concentration LY2109761, 3D cultures derived from AF were exposed to TGFβ or 20 μM LY2109761 in absence or presence of a PD0325901 MEKi (Mirdametinib). Indeed, the MEKi suppressed TGFβ-, as well as 20 μM LY2109761-induced ECM remodeling in a dose-dependent manner with only minimal signs of cytotoxicity at the highest concentrations tested (Fig 7G,H).

In summary, both canonical SMAD signaling and non-canonical ERK MAPK signaling triggered by TGFβ are blocked by low μM concentrations TβRi. This also blocks TGFβ-induced ECM remodeling as expected. Exposure to >10 μM GW788388 or LY2109761 TβRi stimulates ECM remodeling. For LY2109761 we show this occurs in absence of TGFβ and without stimulating canonical SMAD signaling. Lastly, ECM remodeling induced by TGFβ or high concentration LY2109761 is blocked by a MEKi. Together, this indicates that following an early phase in which TGFβ may activate a wider range of signaling mechanisms, non-canonical ERK MAPK signaling is the critical pathway underlying TGFβ-mediated ECM remodeling. It also points to off target effects of high concentrations of the GW788388 and LY2109761 TβRi that hijack the same MEK-mediated pathway, independent of TGFβ.

## DISCUSSION

The transition of fibroblasts to myofibroblasts plays an important role in wound contraction and tissue repair [54]. However, persistence and chronic activation of myofibroblasts drives excessive ECM production and ECM remodeling causing a disorganized tissue architecture, which can ultimately lead to organ failure [55]. There is an urgent need for therapeutic approaches that prevent ongoing fibrosis. Clinical testing of antifibrotic therapeutics includes drugs targeting inflammation or TGFβ signaling [1, 24–27]. Drugs acting further downstream by directly targeting myofibroblast-mediated ECM remodeling may be effective. Yet, targeting myofibroblasts is hindered by a lack of well-defined markers and actionable mechanisms driving fibrotic ECM remodeling are only beginning to be resolved [55–57]. Models that generate quantitative data on 3D fibroblast-mediated ECM remodeling with the reproducibility and throughput needed for drug testing are lacking. The model developed here fits this purpose and produces combined quantitative information on efficacy and cytotoxicity. Our model does not recapitulate the complex interplay between immune cells and fibroblasts that drives *in vivo* fibrosis. However, it does allow the exploration of actionable mechanisms underlying ECM remodeling by the myofibroblasts, which represents a key step in the progression of fibrosis even in the much more complex *in vivo* situation.

The presence of clusters rather than individual, scattered fibroblasts in our model is physiologically relevant as clustering is observed in fibrotic tissue and tissue deformation by myofibroblasts involves interconnected myofibroblast networks [12, 13]. Fibroblasts protruding from the edge of the clusters in our model show aligned actin filaments connecting with extracellular FN fibrils, which, together with the aligned collagen fibers, interconnect fibroblast clusters, thus generating a mechanically coupled network. Formation of this ECM structure requires contractility as evidenced by inhibitors targeting ROCK and LIMK. For inhibitors targeting MLCK, the efficacy largely overlaps with cytotoxicity but targeting non muscle myosin II ATPase with Blebbistatin further demonstrates the potential to suppress ECM remodeling by inhibiting contractility. In agreement, TGFβ-SMAD signaling causes activation of the Rho-ROCK-LIMK pathway [58]. Thus, TGFβ stimulates the expression of FN [38, 39] as well as the contractility required for its assembly into a fibrillar network [59]. In turn, the fibrillar FN network may be important to allow for the collagens produced by TGFβ-stimulated fibroblasts to form a fibrillar network [60] and contractility is also important to align pre-existing collagen fibers between the fibroblast clusters. Moreover, stabilization of this complex ECM structure requires collagen fiber crosslinking since LOXi prevent ECM remodeling as we observe by reflection microscopy. This aligns with studies showing that TGFβ can induce expression of LOX enzymes and implicating some, but not all LOX enzymes in fibrosis [61, 62]. Indeed, LOX inhibition has moved from preclinical to clinical studies for the treatment of patients with lung fibrosis [28](http://clinicaltrials.gov/ct2/show/NCT01242189).

Our findings indicate that S1P can stimulate fibrotic ECM remodeling in a similar, contractility-dependent manner as TGFβ. S1P is a natural bioactive sphingolipid that binds G-protein-coupled receptors and is associated with fibrosis in various tissues such as the liver, lung, and heart [63]. S1P binding to its cellular receptors has been implicated in the activation of ROCK-mediated cell contractility [64, 65]. Given the strong inhibition of cell migration and prominent ECM bridge between fibroblast clusters we observed upon treatment with S1P, its stimulation of Rho-ROCK mediated contractility may be more robust than that achieved with TGFβ. Notably, S1P was only able to stimulate ECM remodeling by clusters of fibroblasts that had been pre-activated with TGFβ (pre-activation with S1P was not effective). Crosstalk between the S1P and TGFβ pathways has been previously reported: TGFβ can enhance S1P synthesis by upregulating sphingosine kinases and a neutralizing S1P antibody reduces TGFβ induced changes in gene expression [66]. One explanation for the requirement for pre-activation with TGFβ for the response to S1P in the fibroblast clusters in our model, is the fact that TGFβ controls expression of cellular S1P receptors [67].

The contractile myofibroblasts that drive ECM remodeling in fibrotic tissues are characterized by several markers of which expression of αSMA is commonly used for their identification [57, 68]. αSMA is induced by a variety of factors including TGFβ but also by mechanical forces applied to the cells [57, 69]. In turn, αSMA supports traction forces applied by fibroblasts onto the ECM [70, 71] pointing to a positive mechanical feedback loop driving formation of contractile myofibroblasts in an αSMA-dependent manner. On the other hand, in mouse models for liver, lung, and kidney fibrosis αSMA positive and negative fibroblasts equally contributed to integrin-mediated contractility-dependent TGFβ activation and production of collagen [72], suggesting that ECM remodeling by activated fibroblasts does not require αSMA.

Our findings indicate that an initial TGFβ mediated transition of fibroblasts to an activated state with expression of αSMA and FN is sustained. I.e., after removal of TGFβ the expression of FN remains high for at least 5 days in AF as compared to NF. However, for ECM remodeling as detected in our model, a continued stimulation by TGFβ is needed and, in this phase, non-canonical TGFβ-mediated activation of the ERK MAPK pathway is required. Activity of the ERK MAPK pathway has been observed by others in fibrosis, including in the liver, lung, and heart [73–75]. In fact, our current work indicates that non-canonical ERK MAPK signaling is the predominant pathway underlying TGFβ-mediated ECM remodeling: First, a MEKi prevents TGFβ-induced ECM remodeling. Second, off target effects of TβRi (see below) stimulate ECM remodeling in absence of TGFβ and they do so in a manner dependent on MEK activity but without any stimulation of canonical SMAD signaling. Constitutive activation of Ras-MAPK signaling caused oncogenic stress in the primary fibroblasts as described earlier [53]. We therefore have not been able to firmly establish that MEK activation is indeed sufficient to promote ECM remodeling in fibroblasts that have already been pre-activated by TGFβ (although we observed a trend in that direction in AF cultures transiently transfected with active MEK resulting in a slight ERK phosphorylation). Nevertheless, our work puts MEK center stage in fibrotic ECM remodeling and the Mirdametinib (PD-0325901) MEK inhibitor that was effective and not cytotoxic in our study may represent a candidate therapeutic approach for fibrosis patients. Interestingly, Mirdametinib is being tested in a Phase 2b clinical trial in patients with neurofibromas and appears to be well tolerated (ClinicalTrials.gov ID NCT03962543).

Given the prominent role of TGFβ signaling in fibrosis and cancer progression the clinical potential of small molecules and antibodies blocking this pathway is extensively explored [76]. However, our findings indicate that the two TβRi we tested may have unanticipated pro-fibrotic effects at concentrations >10 µM. Pharmacological studies are lacking to translate such concentrations to the in vivo situation. Earlier in vitro studies have exposed 2D cultures to 1-20 or up to 100 µM GW788388 or LY2109761 [48, 49, 77–79]. In mouse studies, oral GW788388 or LY2109761 dosing of up to 10 mg/kg/day or 2x50 mg/kg/day, respectively have been reported and one study has reported a GW788388 plasma concentration of 1.5 µM 20 min after oral dosing [80]. Extensive pharmacology to translate this to dynamic exposure in tissues has not been reported. Notably, the decay time of the GW788388 or LY2109761 TβRi in our 3D culture set-up is not known, but over the course of the 72h exposure the concentration is likely to decrease considerably. This probably also explains why <1 µM GW788388 or LY2109761 was ineffective in our study as well as in several previous in vitro studies.

GW788388 and LY2109761 inhibit TβRI/ALK5 with K_i_’s of 18 and 38 nM, respectively (cell-free kinase assays) as well as TβRII, ALK4, and ALK7 with higher K_i_’s [48, 49]. It is unlikely that a shift in the balance of these inhibitory activities as concentrations increase, can explain the apparently pro-fibrotic effect of GW788388 and LY2109761 at higher concentrations. First, TβRII may be more effectively inhibited by these TβRi at higher concentration given the lower K_i_’s, but TβRII has in fact been shown to play a crucial role in TGFβ-induced ERK MAPK activation [81]. Second, increased inhibition of SMAD7 (which mediates a negative feedback loop to TβR signaling) at higher concentrations LY2109761, would be expected to increase canonical TGFβ signaling rather than the selective induction of MEK activity observed by us [82]. Therefore, an off-target effect is likely driving the ECM remodeling appearing at higher concentrations. Indeed, for other ALK inhibitors cell free kinase assays indicate they can target a range of additional kinases causing either decreased or increased activity [83]. It remains to be established which off target is responsible for the enhanced MEK and ROCK dependent ECM remodeling triggered by the two TβRi tested by us, but this effect may raise concern regarding their clinical use.

## MATERIALS AND METHODS

### Cell culture and expression of cDNAs

Primary human dermal and lung fibroblasts were obtained from Lonza and cultured in DMEM Glutamax (31966-021, Gibco, Fisher Scientific, Landsmeer, The Netherlands) supplemented with 10% fetal bovine serum (FBS), 25 U/mL penicillin, and 25 µg/mL streptomycin in a humidified incubator with 5% CO2 at 37 °C. Cells were used between passages 3 and 6.

The lentiviral CAGA_12_-dynGFP construct, expressing superfolder GFP with a destabilizing domain under transcriptional control of twelve 5′-CAGA-3′ SMAD3 response elements, was described previously [84]. The lentiviral pLX311-GFP-MEKDD constitutively active MEK plasmid was a gift from Sefi Rosenbluh obtained through Addgene (plasmid # 194882) [85] CAGA_12_-dynGFP or GFP-MEKDD particles were generated in Lenti-X cells and used to transduce subconfluent fibroblast cultures for 24 hours. Fibroblasts were allowed to recover for 24 hours post-transduction before being used in experiments.

The pHA-MEK^N3;S218E;S222D constitutively active MEK plasmid [86] or a pHA empty vector control plasmid was transfected into NF or AF using Lipofectamine 3000 (ThermoFisher) for 24 hours. Dishes were washed, medium was refreshed, and fibroblasts were allowed to recover for 24 hours post-transfection before being used in experiments.

### Pharmacological inhibitors and antibodies

Pharmacological inhibitors used in this study are listed in Table 1.

Antibodies used for immunofluorescence included mouse anti-human collagen (1:500; PA5-95137, Invitrogen), mouse anti-human FN (1:1000; 610077, BD), mouse anti-human ⍺SMA antibody (1:250; 14-9760-82, Invitrogen), Alexa Fluor 647 goat anti-mouse (1:500; 15-605-146, Jackson), Alexa Fluor 488 goat anti-mouse (1:500; A11001, Invitrogen) or Alexa Fluor 488 goat anti-rabbit (1:500; A11008, Invitrogen).

Antibodies used for Western blot included mouse anti-human ERK (1:1000; 610030, BD Biosciences), rabbit anti-human phospho-Erk1/2 (Thr202/Tyr204) (1:1000; 9101, Cell Signaling), mouse-anti-human FN (1:5000; 610077, BD), mouse anti-human glyceraldehyde 3-phosphate dehydrogenase (GAPDH) (1:1000; sc-32233; Santa Cruz, Dallas, TX, USA), Horseradish peroxidase (HRP)-conjugated goat anti-rabbit IgG (1:10,000; 111-035-003; Jackson Immunoresearch), or HRP-conjugated goat anti-mouse IgG (1:10,000; 115-035-003; Jackson Immunoresearch).

### 3D model for fibrotic ECM remodeling and compound screening

Collagen type I solution was isolated from rat tails by acid extraction as previously described [34, 87]. Collagen (stock concentration: 5 mg/ml) was diluted to 1.5 mg/ml in DMEM containing 0.1 M HEPES (H0887, Sigma-Aldrich) and 44 mM NaHCO3 (stock 440 mM; 71630, Fluka). 70 µl of collagen solution per well was loaded into a 96-well plate (655090, Greiner) and polymerized for 1 hour at 37 °C.

Primary fibroblast cultures were used unstimulated (NF) or activated by exposure to 10 ng/ml TGFβ (TGFβ1; 240-B-002/CF; R&D Systems) for 72 hours (AF). Subsequently, subconfluent monolayers of NF or AF were trypsinized, filtered (04-0042-2317, Sysmex), and re-suspended in 100 µl complete culture medium containing 5% polyvinylpyrrolidone (PVP; P5288, Sigma-Aldrich). Either a single droplet of the PVP/cell suspension per well or multiple droplets per well, spaced 0.9 µm apart, were printed into the collagen gels using an image-guided micro-injection robot (Life Science Methods, Leiden, NL) as previously described [32, 33]. This approach created collagen-embedded clusters of cells with an initial diameter of 200 µm at defined x-y-z positions 150 µm above the bottom of the wells. Subsequently, the 3D cultures were exposed to control medium, 10 ng/ml TGFβ, or 1 µM sphingosine-1 phosphate (S1P) and imaged at several time points over a period of 72 hours. For assays addressing reversibility of ECM remodeling, medium on top of the 3D cultures was subsequently replaced with control medium or medium containing pharmacological inhibitors and monitored for an additional 72 hours.

For compound screening, the 3D cultures were incubated with pharmacological inhibitors 90 minutes prior to adding TGFβ and then monitored over a period of 72 hours.

To evaluate compound cytotoxicity, 1 µg/ml Hoechst33342 (#610959; Thermo Fisher) and 0.4 µM Propidium Iodide (PI) were added to the 3D cultures 1 hour prior to imaging.

### Automated image acquisition and image analysis

All images were acquired using a Nikon TE2000 confocal microscope equipped with a Prior automated stage controlled by NIS Element Software at 20x objective, 20x long distance or 40x long distance water immersion objective in a temperature and CO_2_ controlled incubator.

To analyze ECM remodeling, the collagen fiber network was imaged by confocal reflection microscopy with a 20x or 40x long distance water immersion objective using excitation at 561 nm with a 561 nm blocking dichroic mirror for the detection and capture of the total refection signal. Z-stack images were captured either through a single fibroblast cluster or through two fibroblast clusters, with images taken at 25 μm intervals along the entire z-axis of the cluster(s). To analyze collagen fiber alignment in the single fibroblast cluster model (Fig S1 A,B), CurveAlign v4.0 [37] software was used to obtain a collagen alignment index and the intensity of the aligned collagen fibers. Four 250×250 pixel regions of interest (ROIs) from each corner of every Z-section reflection image were selected and analyzed using the software. The alignment index and intensity of aligned collagen fibers were determined by averaging the values obtained from each ROI in each Z-section. To analyze ECM remodeling in the multiple fibroblast cluster model (Fig 1A,B), reflection microscopy images underwent thresholding to distinguish collagen fibers. Subsequently, a cell-free ROI was defined between 2 clusters using ImageJ. This same ROI was then applied to each Z-section image under all experimental conditions within one biological replicate. The total intensity of the reflection signal in the ROI in each Z-section was determined and averaged. For image processing and batch analysis a macro was created in ImageJ 1.53c.

To assess cytotoxicity of compounds, Hoechst33342/PI-stained 3D cultures were analyzed using confocal fluorescence microscopy with a 20x objective capturing images at 10 µm intervals along the entire z-axis of fibroblasts clusters. Automated image analysis was performed using ImageJ 1.53c and CellProfiler version 2.2.0. In each Z-section, the images from the Hoechst channel were pre-processed by ImageJ to create masked images. Subsequently, Hoechst positive objects were scored as PI positive or negative. The proportion of PI-positive nuclei in each Z-section was determined using an in-house script written by Dr. Joost Willemse (Institute of Biology, Leiden) and numbers for all Z-sections averaged for the entire cluster.

### Immunostaining of ECM embedded fibroblast clusters

Collagen-embedded fibroblast clusters were fixed using 4% paraformaldehyde, washed three times with PBS containing 1% BSA and blocked with PBS supplemented with 1% BSA and 0.3% Triton X-100 for 1 hour at 4°C. Samples were stained with primary antibodies and washed three times with PBS containing 2% BSA, 0.1% Triton X-100, and 0.02% SDS. They were then stained with fluorescently labeled secondary antibodies for 24 hours at 4°C in the dark and washed three times. Lastly, they were stained with 0.05 μM Rhodamine Phalloidin (R415, Thermo Fisher) and 0.4 μg/ml Hoechst 33342 (#610959, Thermo Fisher) for 3 hours at 4°C in the dark and washed three times. Images were captured using a Nikon TE2000 confocal microscope equipped with a long-distance water immersion objective. In the case of CAGA-GFP expressing cells, the average intensity of the GFP signal across the Z-stack was determined using ImageJ 1.53c.

### Western blotting

Cells were lysed with RIPA buffer containing 1% protease/phosphatase inhibitor cocktail (PIC; P8340, Sigma-Aldrich). Samples were separated by SDS-polyacrylamide gel electrophoresis and transferred to polyvinylidene difluoride (PVDF) membranes (Millipore) followed by blocking with 5% BSA in Tris-buffered saline with 0.05% Tween-20. Membranes were incubated with primary antibodies overnight at 4°C followed by HRP-conjugated secondary antibodies for 1 hour at RT. Membranes were developed with enhanced chemiluminescence substrate mixture (ECL plus, Amersham, GE Healthcare, Chicago IL, USA) and imaged using an Amersham Imager (GE, Healthcare Life Science, Chicago, IL, USA).

### Statistical analyses

All statistical analyses were performed in GraphPad Prism 8 using one-way analysis of variance (ANOVA) or two-way ANOVA with Tukey’s post-hoc test. Data were presented as mean and standard error of the mean (SEM). Statistical significance was considered when *P* < 0.05.

## ACKNOWLEDGEMENTS

We thank Joost Willemse (Leiden University) for help with the image analysis.

## Author contributions

E.H.J.D. conceived and supervised the project. C-Y.L., and E.H.J.D. conceptualized and designed experiments. B.C. and P.t.D. provided reagents and critical input in design of experiments. C-Y.L., J.H., and J.C. performed experiments. C-Y.L. and E.H.J.D. wrote the manuscript. All authors reviewed the manuscript and agreed with the final draft.

## Data and materials availability

All data generated or analyzed during this study are included in this published article and its supplementary information files.

