## Supplemental figures for "Novel high throughput 3D ECM remodeling assay identifies MEK as key driver of fibrotic fibroblast activity"

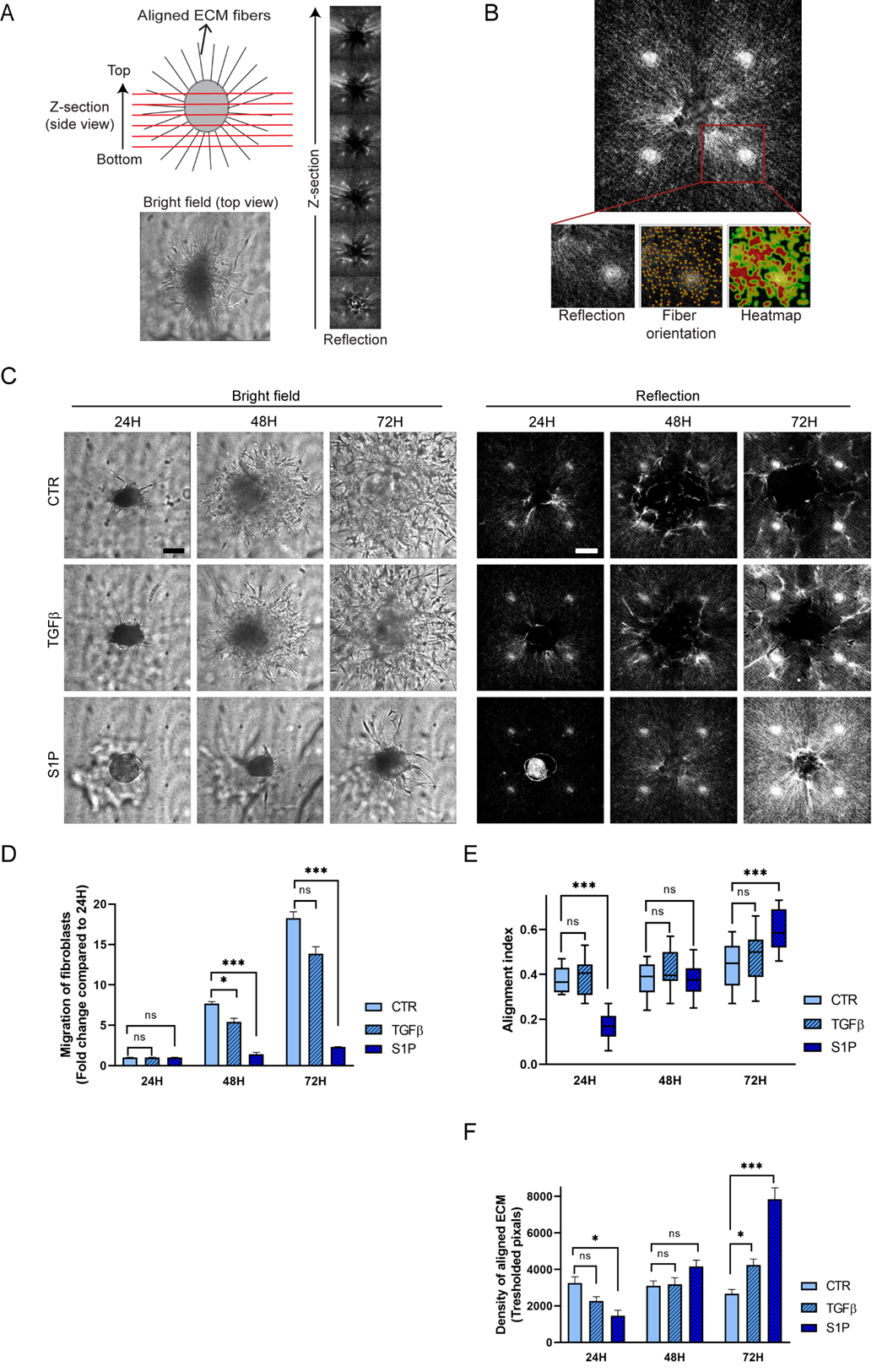

**Figure S1. Single cluster model for quantitative assessment of TGFβ- and S1P-induced ECM remodeling. (A)** Cartoon and brightfield and reflection microscopy images showing aligned ECM fibers perpendicular to the fibroblast cluster measured by reflection microcopy. Z-stack on the right shows reflection images captured at 25 μm vertical spacing throughout the z-axis of a fibroblast cluster. **(B)** Reflection image shows 250 x 250 pixel ROIs in each corner of each Z-section that are selected for analysis using the 'CurveAlign' software (*37*). Each ROI image generates two graphical outputs: fiber orientation and heatmap for calculating the alignment index and quantifying the density of aligned collagen fibers, respectively. Note the white spot in each ROI that is an artefact caused by reflection from optical elements in the microscope (*36*) but which is ignored by CurveAlign as it contains no aligned fibers. **(C)** Brightfield and reflection (single Z-section) images showing outgrowth and ECM remodeling by clusters of AF exposed to TGFβ or S1P for the indicated times. Bar = 100 μm. **(D)** Quantification of fibroblast migration based on brightfield images as shown in (C). **(E,F)** CurveAlign based quantification of reflection signals providing fiber alignment index (E) and the density of aligned fibers (F). An alignment index of 1 indicates a high degree of alignment, while an alignment index of 0 indicates a random distribution of fibers. **(D-F)** Graphs show the mean and SEM of two independent experiments, each performed in duplicate. NS, non-significant; *, p<0.05; ***, p < 0.001.

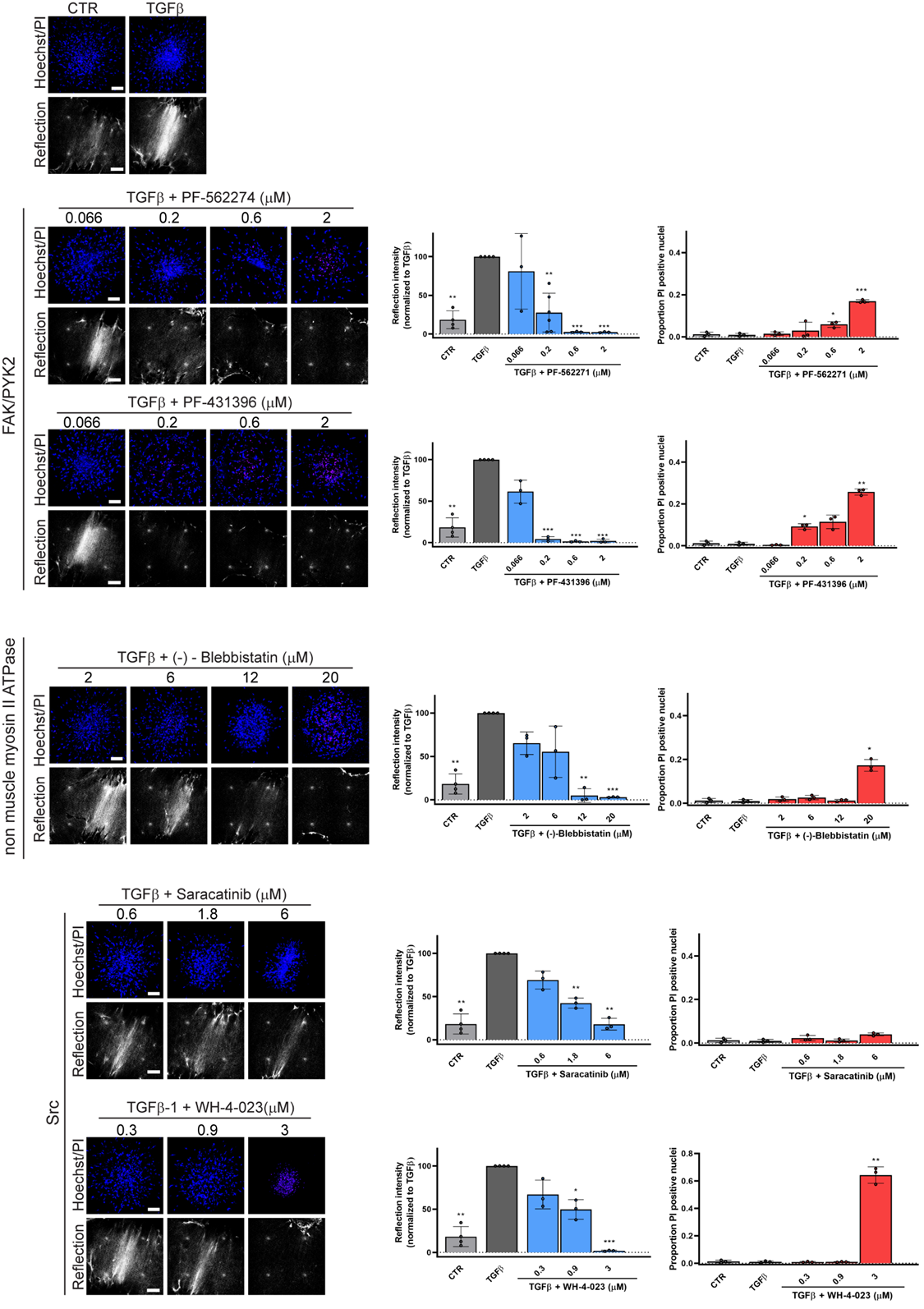

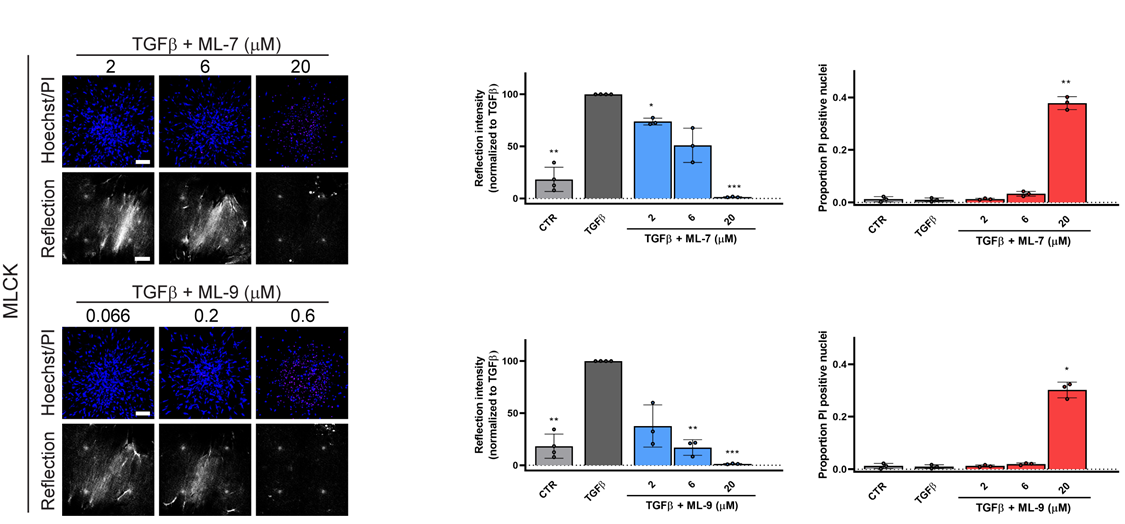

**Figure S2. Overview of the efficacy and cytotoxicity of candidate anti-fibrotic compounds.** Reflection (single Z-section) images of the area between AF clusters and Hoechst (blue)/PI (red) staining (maximum projection) inside AF clusters exposed for 72 hours to TGFβ containing media in absence or presence of the indicated concentrations of the indicated compounds targeting FAK/PYK2, non-muscle myosin II ATPase, Src, and MLCK. Bar = 100 μm. Graphs show quantification of reflection signals and Hoechst/PI staining. Mean and SEM of two independent experiments, each performed in triplicate is shown. *, p < 0.05; **, p < 0.01; ***, p < 0.001 compared to TGFβ alone condition for reflection and compared to CTR condition for PI staining.

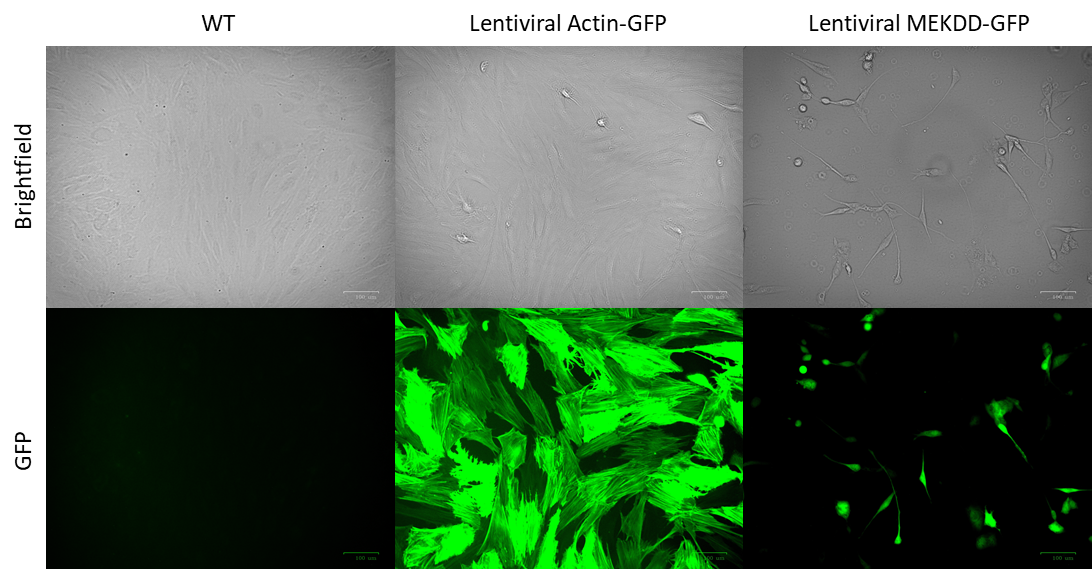

**Figure S3. Lentiviral expression of constitutively active MEK leads to cell death in primary human fibroblasts.** Brightfield and GFP fluorescence microscopy for primary human fibroblast cultures under control condition of transduced with lentiviral particles expressing the indicated constructs.
